# Mitochondrial Raf1 Regulates Glutamine Catabolism

**DOI:** 10.1101/2024.03.08.581297

**Authors:** Ronald L. Shanderson, Ian D. Ferguson, Zurab Siprashvili, Luca Ducoli, Albert M. Li, Weili Miao, Suhas Srinivasan, Mary Grace Velasco, Yang Li, Jiangbin Ye, Paul Khavari

## Abstract

Raf1 is present within the mitochondrial matrix, where it binds GLS to regulate glutamine catabolism and tumorigenesis.

In cancer, Raf1 activation occurs via mechanisms that include mutation of upstream regulators, such as receptor tyrosine kinases and Ras GTPases, as well as by mutations that affect *RAF1* itself, including via gene amplification (*1*–*4*). Once recruited to the plasma membrane (**<u>PM</u>**) Raf1 can engage downstream mitogen-activated protein kinase (**<u>MAPK</u>**) pathway signaling through phosphorylation of the MEK kinases (*5*). In addition to Raf1, A-Raf and B-Raf can also activate MEK and these other two Raf isoforms can compensate for MAPK activation in the event of Raf1 loss (*6*, *7*). Despite this, Raf1 remains essential for the development and maintenance of some tumors through mechanisms independent of MAPK activity (*7*, *8*). In this regard, Raf1 has well-described interactions outside the canonical MAPK pathway, including several with outer mitochondrial membrane (**<u>OMM</u>**) proteins (*9*, *10*), although Raf1 has not been previously identified inside mitochondria. Mitochondria comprise a hub for various metabolic processes modulated in cancer cells to accommodate rapid proliferation. One such process is glutaminolysis, which involves the catabolism of glutamine to generate both ATP as well as precursors for the synthesis of fatty acids, nucleotides, and nonessential amino acids (*11*–*13*). Glutaminase (**<u>GLS</u>**) proteins, which catalyze the first and rate-limiting step of this process by converting glutamine to glutamate, are often upregulated in cancer (*14*–*16*). GLS activation has been previously associated with tumors driven by Ras, upstream regulators of Raf kinases (*13*, *17*). Here we identify Raf1 protein inside mitochondria where Raf1 associates with GLS in the mitochondrial matrix to enable glutamine catabolism and tumorigenic growth.

Raf kinases play vital roles in normal mitogenic signaling and cancer, however, the identities of functionally important Raf-proximal proteins throughout the cell are not fully known. Raf1 proximity proteomics/BioID in Raf1-dependent cancer cells unexpectedly identified Raf1-adjacent proteins known to reside in the mitochondrial matrix. Inner-mitochondrial localization of Raf1 was confirmed by mitochondrial purification and super-resolution microscopy. Inside mitochondria, Raf1 associated with glutaminase (GLS) in diverse human cancers and enabled glutaminolysis, an important source of biosynthetic precursors in cancer. These impacts required Raf1 kinase activity and were independent of canonical MAP kinase pathway signaling. Kinase-dead mitochondrial matrix-localized Raf1 impaired glutaminolysis and tumorigenesis in vivo. These data indicate that Raf1 localizes inside mitochondria where it interacts with GLS to engage glutamine catabolism and support tumorigenesis.

## Results

### Raf1 proximal proteins nominated by BioID

To identify Raf1-adjacent proteins of interest in living cancer cells, the BASU promiscuous biotin ligase was fused to Raf1, then full-length fusion protein was expressed at endogenous levels and biotin labeling activity verified (**Fig. 1A, fig. S1A-D**) (*18*, *19*). Raf1-BASU, but not eGFP-BASU fusion control, labeled positive control MEK MAPK proteins, demonstrating that the Raf1 fusion retains proximity to known physiologic interactors (**fig. S1B**). Raf1-BASU was then expressed in Raf1-dependent MM485 (melanoma) and AsPC1 (pancreas) cancer cell lines as well as matched Raf1-independent CHL1 and BxPC3 cancer cell lines derived from the same tumor types (*20*), namely melanoma and pancreatic cancer, respectively (**fig. S1E**). Biotin labeling followed by streptavidin pull-down and LC-MS/MS was then performed (*18*, *19*). This nominated a number of proteins as proximal to Raf1, including 82 previously-identified Raf1 interactors, including Mek1, Mek2, YWHAQ, YWHAZ, B-Raf, and SPRY4 (**Fig. 1B, table S1**) (*21*). These findings indicate that this BioID approach can identify biologically relevant Raf1-associated proteins.

**Fig. 1.**
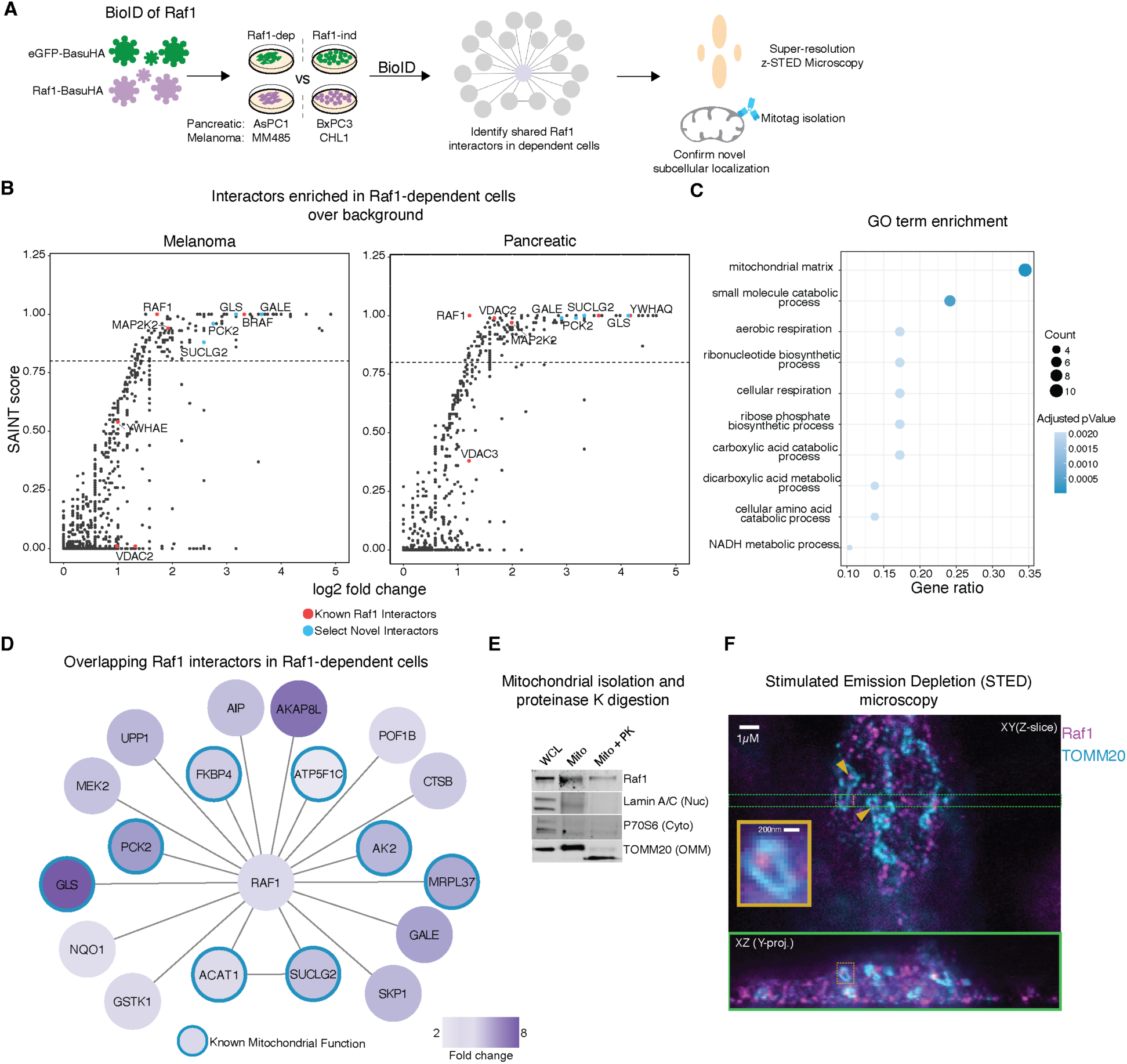
Raf1 proteomics reveals mitochondrial localization. **(A)** Schematic of Raf1 BioID and localization workflow. (**B**) SAINT plots demonstrating all hits with a positive SAINT score for proteins enriched in the proximal proteome of Raf1 in dependent MM485 or AsPC1 cells compared to Raf1-independent CHL1 and BxPC3 and eGFP controls. Red are select known Raf1 interactors and cyan indicates proteins of interest involved in metabolic processes. Dashed line drawn at SAINT score of 0.8 (**C**) Combined Cell Component, Biological Process, and Molecular Function Gene Ontology Enrichment in SAINT ≥ 0.8 proteins. Benjamini-Hochberg adjusted pValues are used. (**D**) Network of Raf1 proximal proteins with SAINT ≥ 0.9 in both pancreatic and melanoma cell lines of interest. Edges between non-Raf1 proteins represent known interactions. Proteins with known mitochondrial localization labeled with blue. (**E**) Mito-tag Mitochondrial isolation of MM485 melanoma cells. WCL denotes whole cell lysate, Mito indicates mitochondrial fraction, and PK denotes proteinase K treatment to digest outer mitochondrial membrane (OMM) proteins. Lamin A/C is a nuclear (nuc) protein, p70S6 kinase is a cytoplasmic (cyto) protein, and TOMM20 spans the mitochondrial outer mitochondrial membrane (OMM) (**F**) STED microscopy demonstrating inner-mitochondrial Raf1 (yellow arrows). Image stack side-view (XZ and YZ) are cropped laterally and then mean-projected along X or Y. Inset displays a Y-projection of a particular mitochondria of interest in XZ.

This dataset also included a number of novel putative Raf1-proximal proteins. Most notably, gene ontology analysis of interactors associated with Raf1 in Raf1-dependent cells revealed enrichment for proteins associated with the mitochondrial matrix (**Fig. 1C**, **table S2**), a subcellular location where Raf1 had not previously been described. In Raf1-dependent cancer cell lines, 38% of the most highly enriched Raf1-proximal proteins are known to localized inside mitochondria, among which is GLS (**Fig. 1D**) (*14*); such an enrichment was not seen in Raf1-independent cancer lines. This finding suggested a previously undescribed localization for Raf1 inside mitochondria in certain contexts.

To explore this possibility, mitochondria were purified then treated with proteinase K to remove outer mitochondrial proteins. Digestion of the extra-mitochondrial domain of the TOMM20 trans-membrane protein verified removal of proteins on the OMM; proteins inside mitochondria were retained, as demonstrated by detection of the inner-mitochondrial portion of TOMM20 (*22*). The Raf1-BASU fusion was observed in Raf1-dependent MM485 melanoma cells, but not in Raf1-independent CHL1 melanoma cells (**fig. S1F**). To further verify the localization of endogenous Raf1, mitochondria were isolated using the mito-tag protocol (*23*). Purified mitochondria treated with proteinase K confirmed an inner-mitochondrial localization of the endogenous Raf1 protein (**Fig. 1E**, **fig. S1G**). For orthogonal validation of Raf1 localization inside mitochondria, super-resolution stimulated emission depletion (**<u>STED</u>**) microscopy was performed using optical sectioning allowed by z-STED (**Fig. 1F**), confirming Raf1 localization inside mitochondria.

### Raf1 impacts on glutaminase and MAPK pathway signaling

In addition to GLS, PCK2 and SUCLG2 were also among the mitochondrial proteins found to be proximal to Raf1 by BioID. The known functions of these mitochondrial matrix-localized proteins suggested that mitochondria-localized Raf1 might influence glutamine catabolism and the citric acid cycle (**<u>TCA</u>**). To explore this, ^13^C-glutamine tracing was performed with and without Raf1 depletion (**Fig. 2A**); MM485 was compared to CHL1 melanoma cells because BioID identified a stronger Raf1 mitochondrial localization in the former. Because GLS is the first and rate-limiting step of glutaminolysis (*24*), GLS depletion was also performed in parallel (**fig. S2A**), as a positive control to help benchmark the point at which glutaminolysis might be influenced by Raf1 activity. After glutamine starvation, labeled glutamine was reintroduced and metabolites were measured before steady state metabolite labeling occurred. Consistent labeling of glutamine was observed in both cell lines across all conditions, suggesting that glutamine uptake was unaffected by Raf1 loss in this setting (**Fig. 2B**, **table S3**). Loss of Raf1 reduced the labeling fraction (M+5) of glutamine-derived-glutamate as well as alpha-ketoglutarate in Raf1-dependent MM485 cells that contained Raf1 localized within mitochondria (**Fig. 2B**). This was also true for 2-hydroxyglutarate, which is derived from alpha-ketoglutarate (**fig. S2C**) (*25*). Further, labeled citrate demonstrated that Raf1 loss alters glutamine that is being processed reductively (M+5) as well as oxidatively (M+4) (**Fig. 2C**) (*26*). Raf1 has been previously linked to glutathione-S transferase P1 (GSTP1), however, no significant changes in the amount of labeled reduced glutathione were observed with Raf1 loss in MM485 cells (**fig. S2D**) (*27*). These data suggest that mitochondria-localized Raf1 may modulate the TCA at the level of GLS.

**Fig. 2.**
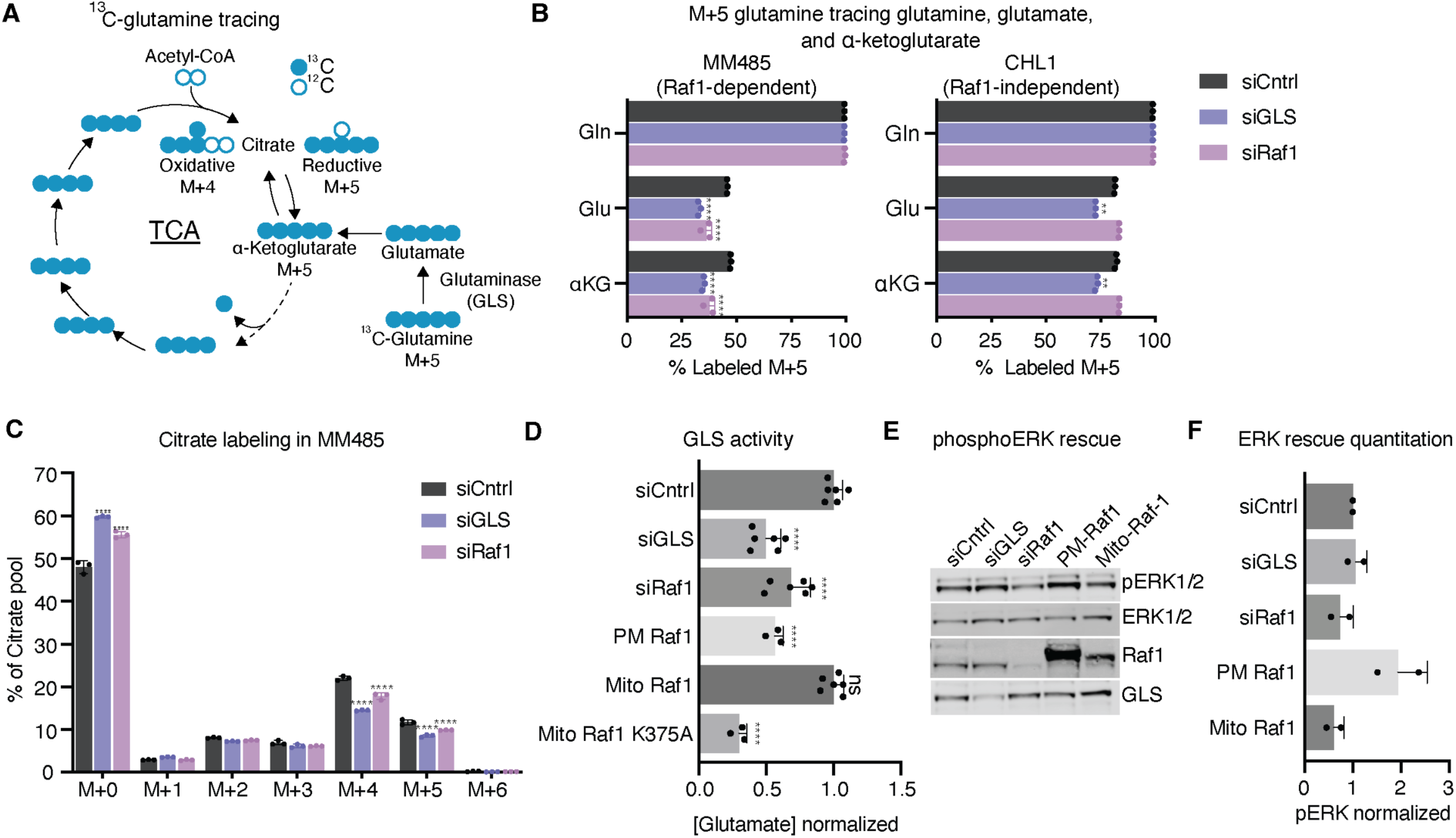
Mitochondrial Raf1 regulates glutaminolysis. **(A)** Schematic indicating ^13^C glutamine tracing for both reductive and oxidative TCA. (**B**) 13C-Glutamine tracing of M+5 Glutamine (Gln), Glutamate (Glu), and αKetoglutarate (αKG) with siRNA knockdown of GLS or Raf1; n = 3, **** *P* < 0.0001, ** *P* < 0.01. (**C**) Mass isotopologues of citrate pool in MM485 cells; n = 3,\*\*\*\**P* < 0.0001. (**D**) Glutamine to glutamate conversion as measured by luciferase-based Glutamine Glo assay with a non-targeting siRNA as a negative control. All Raf1 knockdowns rescued with differentially localized proteins. siCntrl (n =6) is nontargeting siRNA, siGLS (n=6), siRaf1 (n=6), and rescue with PM Raf1 (n=3), Mito Raf1 (n=6), or Mito Raf1 K375A kinase dead construct (n=3); **** *P* < 0.0001. (**E**) Western blot of MM485 cells blotting for ERK activation, effective Raf1 knockdown, Raf1 rescue, and successful GLS knockdown.(**F**) Quantitation of 2 western blots with phosphoERK normalized to total ERK.

To study if Raf1 localization inside mitochondria may be necessary for the observed effects on glutaminolysis, differentially-localized Raf1 constructs were produced. A mitochondrial matrix-localized Raf1 (**Mito Raf1**) was generated by fusing the protein to the mitochondrial-localized domain of COX4I1. As a control, a plasma membrane-localized Raf1 (**PM Raf1**) was created by adding the membrane-localization domain of the XRP2 protein to Raf1. Previous work demonstrated that PM Raf1 induces MAPK pathway activity (*28*). To assess Raf1 impacts on glutaminase activity, the conversion of glutamine to glutamate (*29*) was quantified. Congruent with glutamine tracing data, the amount of glutamine converted to glutamate decreased with Raf1 loss (**Fig. 2D**). Although it failed to activate the MAPK pathway, Mito Raf1 rescued the loss glutaminase activity, however, PM Raf1 did not (**Fig. 2D**), even though PM Raf1 induced the MAPK pathway (**Fig. 2E-F**). Mitochondrial delivery of the Raf1 K375A mutant, which alters Raf1 structure and abolishes Raf1 kinase activity (*28*), produced lower glutaminase activity than loss of Raf1 alone, suggesting a possible dominant-negative effect (**Fig. 2D**). These findings suggest that Raf1 effects on glutaminase activity and MAPK signaling are separable, and that mitochondrial-localized Raf1 mediates the former.

### Raf1 association with GLS

The observations that Raf1 modulates glutaminase activity, that it can be detected in the mitochondrial matrix where GLS is canonically localized (*14*), and that it is proximal to GLS by BioID raised the possibility that Raf1 might interact physically with GLS. Consistent with Raf1 proximity to GLS, proximity ligation assay (**<u>PLA</u>**) in MM485 cells detected a Raf1-GLS signal; this signal was specific as it was diminished by Raf1 and GLS knockdown (**Fig. 3A, fig. S3A-C**). Additionally, Raf1 and GLS also displayed reciprocal co-immunoprecipitation; this was selective in that neither Ras nor MEK proteins brought down GLS (**Fig. 3B, fig. S3D**).

**Fig. 3.**
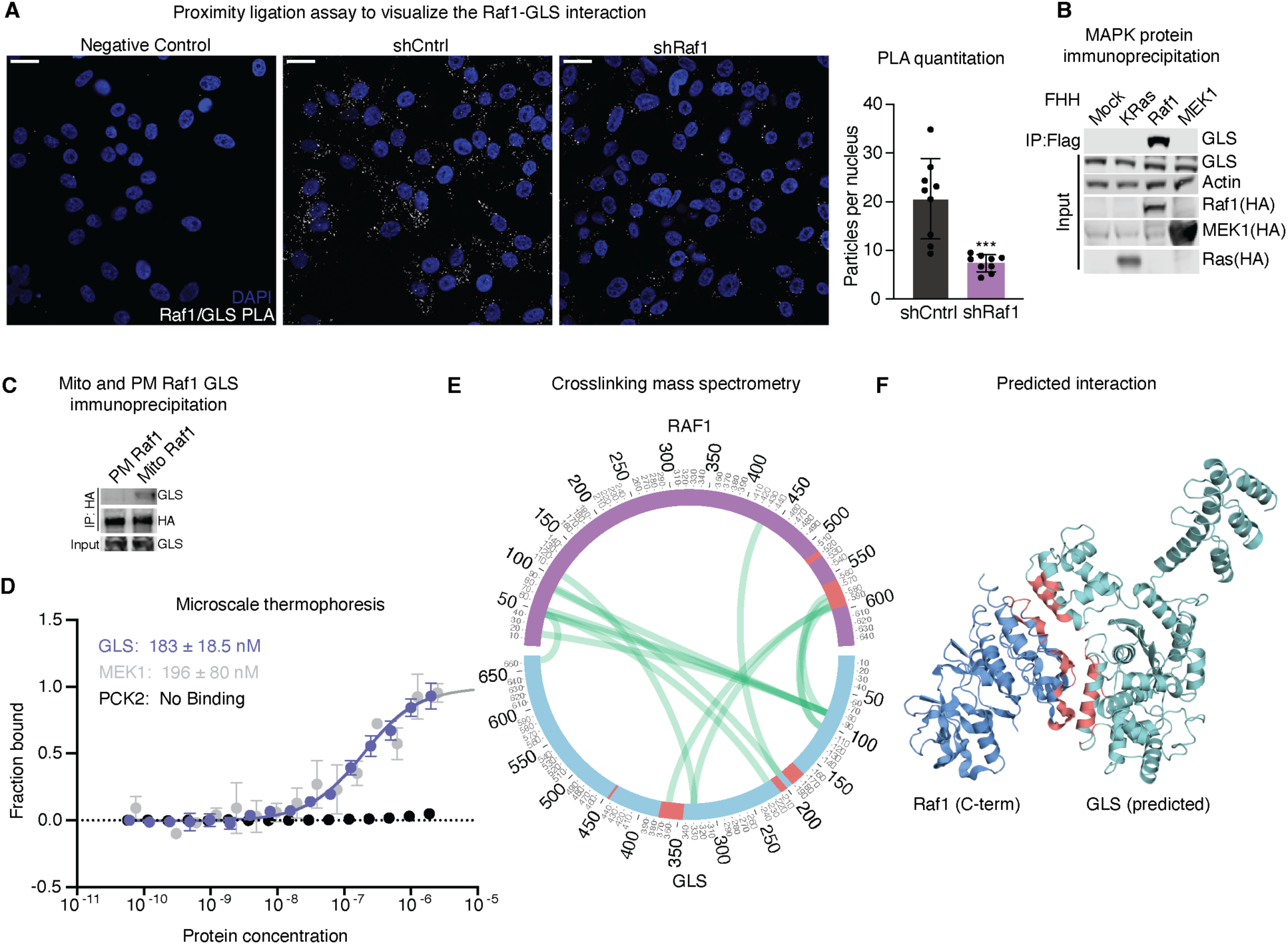
Raf1 directly interacts with GLS. **(A)** PLA between Raf1 and GLS with shRNA knockdown of Raf1 with control. Scale bar represents 20 µm. Includes quantitation in bar graphs quantify particles per nucleus; *** *P* < 0.001 **(B)** Co-immunoprecipitation of Flag-6XHis-HA tagged MAPK component proteins with a flag antibody with appropriate inputs. (**C**) co-IP of plasma membrane and mitochondrial Raf1 with immunoblot for GLS with appropriate inputs. (**D**) Microscale thermophoresis of labeled 6X His-Raf1 against GLS, MEK1, or PCK2 to produce affinity constants of 183 nM, 196 nM, and no binding respectively. (**E**) Crosslinking mass spectrometry circos plot between crosslinked GLS and Raf1. Pink indicates the interface between Raf1 and GLS according to docking experiments. **(A)** (**F**) Molecular docking between Raf1 in blue and GLS in green. Interfacing amino acids indicated in salmon.

Consistent with a mitochondrial location for Raf1-GLS interactions, Mito Raf1 robustly immunoprecipitated GLS whereas PM Raf1 did not (**Fig. 3C**). To examine the possibility that Raf1 and GLS proteins bind each other directly, purified recombinant proteins were studied by microscale thermophoresis (**<u>MST</u>**) (**fig. S3E**). Raf1 displayed a binding affinity to GLS (*K_d_*=1.83x10^-7^M) comparable to its affinity for Mek1 (*K_d_*=1.96x10^-7^M) (**Fig. 3D**). In contrast, another mitochondrial protein detected by Raf1 BioID, namely PCK2, failed to bind Raf1 (**Fig. 3D**), suggesting that associations between PCK2 and Raf1 in the mitochondrial matrix may be indirect. These data suggest that Raf1 can bind GLS directly at affinities comparable to well-characterized Raf1-interacting proteins.

To gain additional insight into the Raf1 association with GLS, interacting regions of Raf1 and GLS proteins were mapped. First, crosslinking mass spectrometry (**<u>CLMS</u>**) was used to identify points of interaction between Raf1 and GLS. Purified recombinant Raf1 and GLS protein were crosslinked to one another using bis(sulfosuccinimidyl)suberate (**<u>BS3</u>**) (**fig. S3F**) then mass spectrometry was performed. A number of crosslinked peptides were identified between Raf1 and GLS, with prominent signal between the N-terminal half of GLS and specific C- and N-terminal portions Raf1 (**Fig. 3E, fig. S3G**). Molecular docking simulations agreed with CLMS data (**Fig. 3F**), further supporting the existence of multiple Raf1-GLS contact interfaces in those protein regions.

### Mitochondrial Raf1 in experimental tumorigenesis and spontaneous human cancers

The impact of mitochondria-localized Raf1 was next studied in experimental tumorigenesis. Wild-type Raf, PM Raf1, Mito Raf1, and Mito Raf1 K375A kinase were expressed in MM485 melanoma cells (**fig. S4A**). After subcutaneous injection in immune deficient mice, cells with enforced expression of wild-type and mito Raf1 displayed similar tumorigenic growth in vivo that were increased over PM Raf1 (**Fig. 4A**). In contrast, Mito Raf1 K375A expressing tumors grew significantly more slowly than others (**Fig. 4A**). Notably, Mito Raf1 K375A did not decrease ERK phosphorylation compared to cells not expressing the construct, suggesting that this abrogation of growth was not due to negative effects on global MAPK activity (**Fig. 4B-C**). Differences in tumorigenic growth in vivo were not reflected in proliferation in vitro (**fig. S4B**), indicating these impacts were not due to global impacts that alter cell viability or capacity for growth. To assess if the Raf1-GLS interaction occurs in spontaneous human malignancies, PLA was performed on a series of 26 spontaneous epidermal squamous cell carcinomas (**<u>SCC</u>**) (*30*), which are associated with Ras-MAPK activation, along with 6 independent normal skin controls. Epithelial cells in SCC displayed substantially more Raf1-GLS signal compared to normal epidermis on a per cell basis (**Fig. 4D-E, fig. S5A-B**). Raf1-GLS PLA on 42 additional human cancer specimens from breast, bladder, ovary, liver, pancreas, and prostate detected a range of Raf1-GLS PLA signals (**fig. S5D-E**). Taken together, these findings indicate that mitochondrial Raf1 can influence experimental tumorigenesis and that Raf1-GLS adjacency can be detected in a subset of spontaneous human tumors.

**Fig. 4.**
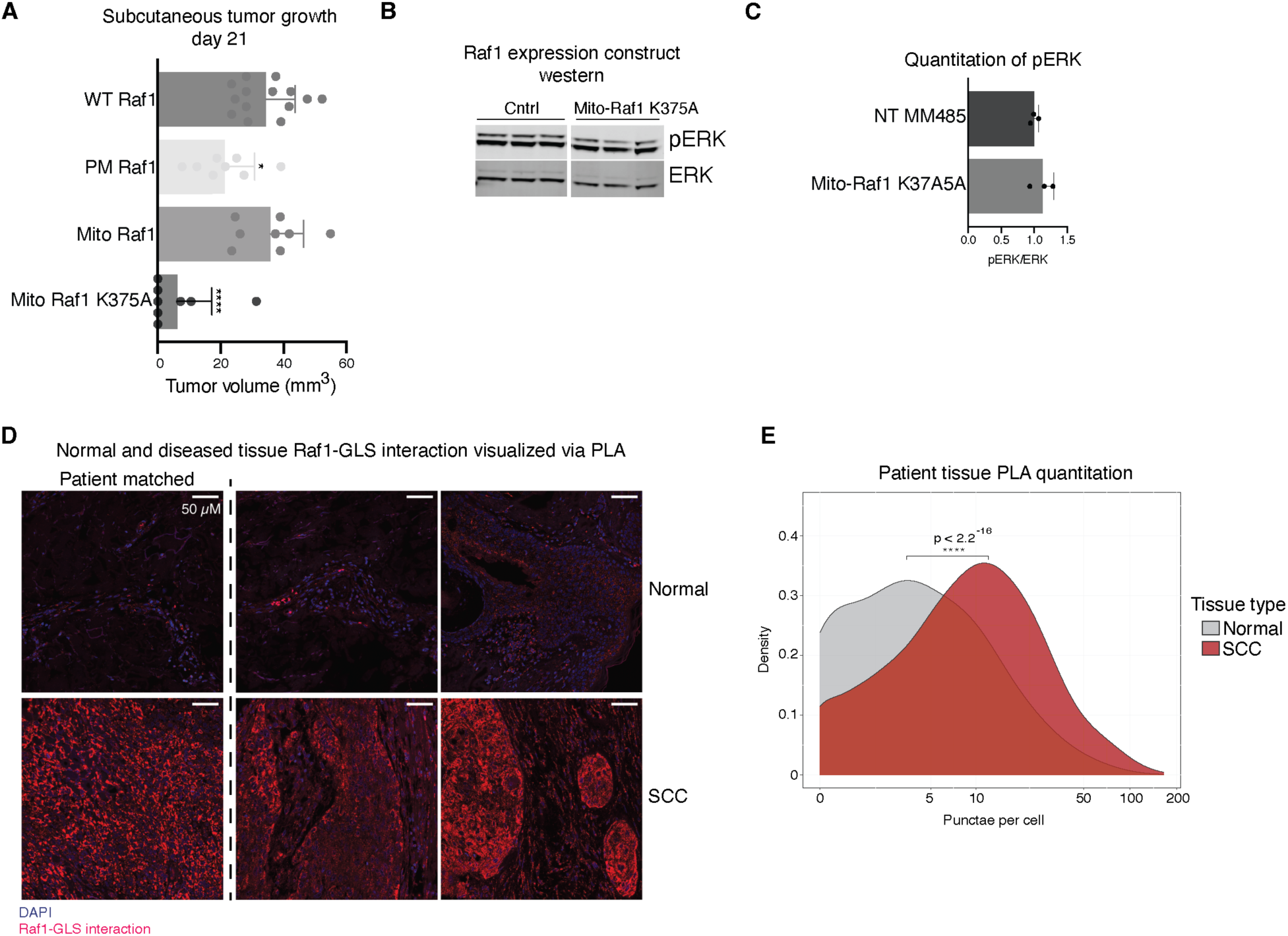
Mitochondrial Raf1 and GLS interaction contributes to tumorigenesis and is present in patient tumors. **(A)** Subcutaneous tumor growth with overexpression of 4 Raf1 constructs: WT Raf1 (n = 14), PM Raf1 (n = 8), Mito Raf1 (n = 8), and Mito Raf1 Kinase Dead (K375A) (n = 8); * *P* < 0.05, **** *P* < 0.0001. (**B**) Western blot of phosphorylated ERK and total ERK protein in cells expressing empty vector or mitochondrial Raf1 K375A. (**C**) With quantitation; *P* = ns. (**D**) PLA of spontaneous tumor microarrays for 3 normal and 3 squamous cell carcinoma (SCC) samples, including one matched. Scale bar represents 50 µm. (**E**) Density plot of puncta count per cell for PLA across several patient samples: 6 normal skin samples, and 26 SCC samples. Mean puncta count per cell is 6.928 for normal skin and 14.266 for SCC; **** *P* < 0.0001.

## Discussion

These studies identified Raf1 inside mitochondria in certain contexts where it is proximal to a variety of proteins native to the mitochondrial matrix, including GLS. The mechanisms responsible for Raf1 translocation into mitochondria remain to be explored. Previous work, however, demonstrated HSP90-dependent mitochondrial translocation of other kinases, such as Akt1 (*31*). Mitochondria-localized Raf1 enables glutaminase activity and supports tumorigenesis without substantial impacts on MAPK signaling. This points to a non-canonical role for Raf1 in supporting glutaminolysis, a process which contributes biosynthetic precursors and energy in neoplasia. Impaired glutamine catabolism seen with global Raf1 loss can be rescued by mitochondrial-localized Raf1 but not Raf1 targeted to the plasma membrane, highlighting the potential role that Raf1 subcellular localization plays in this setting. The K375A Raf1 point mutation, which alters Raf1 protein conformation and kinase activity, disrupts both glutamine catabolism and tumorigenesis, underscoring the importance of the intactness of this Raf1 region in these effects. We were unable, however, to obtain evidence that GLS is a direct phosphorylation target of Raf1, providing a rationale for future efforts to define how Raf1 may modulate GLS activity. Recombinant Raf1 and GLS proteins associate directly at an affinity comparable to Raf1 binding to its well-studied downstream target, Mek1. CLMS and structure modeling nominated specific interface regions between GLS and Raf1. Evidence for Raf1-GLS proximity was observed in tumors from variety of tissues, suggesting that this interaction may play a potential role in spontaneous human cancers.

## Acknowledgments

We thank LV Jackrazi and RT Brennan for experimental help, AES Barentine for STED visualization assistance, and G Kim for an Image J script for processing PLA images. This work was supported by AR045192 and AR049737 from NIAMS/NIH to PAK and also in part by NIH P30 CA124435 for the Stanford Cancer Institute Proteomics Shared Resource. Jiangbin Ye is a Stanford Maternal and Child Health Research Institute Research Scholar. We would also like to thank PK Jackson, JE Ferrell, and JZ Long for prereview.

## Author contributions

Conceptualization: RLS, PAK

Methodology: RLS, IF, ZS, AL, PAK

Investigation: RLS, IF, ZS, LD, AL, WM, SS,YL

Visualization: RLS, SS

Formal Analysis: RLS, SS

Funding acquisition: PAK, JY, RLS

Project administration: RLS, PAK

Supervision: PAK, JY

Writing – original draft: RLS, PAK

Writing – review & editing: RLS,LD, PAK

Competing interests:

Authors declare that they have no competing interests

## Materials and Methods

### Cell culture

Cells were grown in appropriate media containing 10% vol/vol fetal bovine serum (FBS), 100 U/ml penicillin, 100 µg/ml streptomycin, and 250 ng/ml of Gibco Amphotericin B. Briefly, the following cell types were used with appropriate media in parentheses: MM485 (RPMI), CHL1(DMEM), AsPC1 (RPMI), BxPC3 (RPMI).

### Viral transduction

All exogenous expression constructs were cloned into either LentiORF pLEX-MCS-IRES-Puro (OHS4735, Open Biosystems) or pBABE-puro (Plasmid #17643, addgene). For shRNA constructs, PLKO.1 (Plasmid #8453, Addgene) backbone was used. The BASU sequence was previously published (Ramanathan et al. *Nature Methods*, 2018). Raf1 construct was cloned from pLEX constructs from existing lab stocks, though modifications were made to create siRNA and shRNA resistant constructs via inverse PCR.

### Protein-BasuHA fusion

Raf-1 and eGFP fusion proteins to Basu was created by fusing the Raf1 or eGFP CDS to that of a BASU biotin ligase (*18*) on the c-terminal end of the protein because previous data on N-terminal fusions suggest it prevents Raf1 from interacting with Ras appropriately. The proteins are connected with a Glycine/Serine linker with the amino acid sequence ‘Gly-Ser-Gly-Gly-Gly-Ser-Gly-Gly-Gly-Ser’. The BASU was directly fused to an HA tag, ‘Tyr-Pro-Tyr-Asp-Val-Po-Asp-Tyr-Ala’

### Immunoblotting

Cells were lysed in 1X RIPA lysis buffer with Complete Mini Protease Inhibitor Cocktail, EDTA-free (Millipore Sigma) PhosStop phosphatase inhibitor cocktail (Millipore Sigma). Lysates were sonicated at 10% amplitude for 10 seconds and centrifuged at 16,000 x g for 12 minutes. Soluble fraction was removed and quantified using Pierce BCA Protein Assay Kit. 15-25 µg of protein was loaded into a 4-12% NuPAGE novex Bis-Tris gradient gel (Thermo Fisher Scientific) and ran at 170 Volts for 1 hour or until ladder was appropriately resolved. Proteins then transferred to 0.45 µM Immobilon-FL PVDF (Millipore Sigma) after methanol activation. Protein-containing-membranes blocked with LI-COR Odyssey blocking buffer (TBS) for 1 hour at room temperature before being treated with primary antibodies diluted in Odyssey blocking buffer overnight at 4°C. After primary incubation, membranes washed 3X 8 minutes in TBST. Secondary antibodies were diluted 1:20,000 in 5% nonfat milk and 0.02% SDS in TBST and incubated for 1 hour at room temperature. Membranes again washed 3X 8 minutes in TBST and a final 5 minute wash in PBS was conducted before imaging on the Odyssey CLx. All immunoblot quantification done in Image Studio Lite.

Primary antibodies used: p44/42 (ERK1/2)(L34F12)(4696S, Cell Signaling Technology), Phospho-p44/42 MAPK (ERK1/2)(Thr202/204)(9101S, Cell Signaling Technology),c-Raf (D4B3J) (537455, Cell Signaling Technology), Tom20 (D8T4N)(424065, Cell Signaling Technology), p70 S6 Kinase (49D7)(2708S, Cell Signaling Technology), Lamin A/C (4C11)(4777S, Cell Signaling Technology), beta-tubulin(9F3)(2128S, Cell Signaling Technology), SUCLG2(C-1)(sc-393756), HA-Tag (C29F4, Cell Signaling Technology), Glutaminase[EP7212] (ab156876, abcam) Secondary antibodies used: IRDye 800CW Goat anti-Mouse IgG, IRDye 680RD Goat anti-Mouse IgG, IRDye 800CW Goat anti-Rabbit, IRDye 680RD Goat anti-Rabbit

### Proximity-dependent biotin labeling

MM485, CHL1, BxPC3, AsPC1, HT1376, or expressing either pLEX eGFP-BASU-HA or pLEX Raf1-BASU-HA were grown to 85% confluence in 1 15cm plate per replicate. Cells were starved of biotin overnight before labeling. Cells were labeled with 50 µM biotin in appropriate media for 4 hours. After labeling, cells were washed with PBS before being incubated in 15 ml of DPBS at 4°C for 15 minutes to allow for excess biotin to diffuse out of the cells. Cells then removed using 0.05% trypsin-EDTA (25300054, ThermoFisher). Trypsin quenched with 10% FBS-containing-media and pelleted at 1000 x g for 5 minutes. Pellets were lysed in 325 µl of BioID lysis buffer (50 mM Tris pH 7.4, 0.5 M NaCl, 0.2% SDS, 1mM DTT in H2O). After lysis, 28.25µl of 25% Triton X-100 was added to prevent SDS precipitation and samples were placed on ice for sonication at 10% amplitude for 10 seconds. Samples were diluted with 353 µl of cold 50 mM Tris pH 7.4 before repeating sonication. Lysates cleared at 16,000 x g at 4°C for 15 minutes. Biotinylated proteins were extracted using as previously described (Roux et al. 2012) with a few changes: KingFisher Flex Sample Purification System was used at 4°C to perform binding and washes, 2% Sodium Dodecyl Sulfate was replaced with 2% Lithium Dodecyl Sulfate to prevent precipitation in 4°C for wash buffer 2, and MagReSyn Streptavidin beads (MR-STV005, Allumiqs) were used to isolate biotinylated proteins. Washed beads resuspended in TEAB and stored at 4°C before mass spectrometry sample prep. A portion of beads had biotinylated proteins extracted with 1X NuPAGE LDS Sample Buffer, 20 mM DTT, 4 mM biotin and was run along with input and unbound to confirm effective pulldown and transferred to a PVDF membrane. Membrane was probed with IRDye 800CW Streptavidin (P/N: 926-32230, Li-COR).

### Mass spectrometry sample prep

Beads resuspended in 200 µl of 100 mM TEAB buffer before reduction in 10 mM DTT (10197777001, MilliporeSigma). DTT activated on heat block for 5 minutes at 55°C. Samples then mixed head-over-head for 30 minutes. After reduction, samples alkylated with 30 mM acrylamide and mixed for 30 minutes. 0.5 µg of trypsin/Lys-C mixture (V5073, Promega) added to each on-bead sample and set to rotate head-over-head overnight at RT. Trypsin quenched with 1.5% MS grade formic acid (94318, Honeywell). Acidified peptides quantified using Qubit 4 Fluorometer (ThermoFisher). All following spin steps occur at 1,600 x g unless otherwise noted. C18 MonoSpin column (5010-21701, GL Sciences Inc) equilibrated with 200 µl of 50% LC-MS Grade acetonitrile(047138.K2, ThermoFisher) spun through for 1 minute. Equilibrated column washed with 2x 200 µl for 1 minute with 0.1% formic acid. Acidified and quantified peptides spun through C18 column at 1600 x g for 3 minutes, flow through was rebound and spin was repeated. Columns were again washed 2x with 200 µl 0.1% formic acid.

### Peptide mass spectrometry

Peptide pools were reconstituted and injected onto a C18 reversed phase analytical column, ∼25 cm in length packed in house using Reprosil Pur. The UPLC was a Waters NanoAcquity, operated at 450nL/min using a linear gradient from 4% mobile phase B to 35% B. Mobile phase A consisted of 0.1% formic acid, water, Mobile phase B was 0.1% formic acid, water. The mass spectrometer was an Orbitrap Elite set to acquire data in a data dependent fashion selecting and fragmenting the 15 most intense precursor ions in the ion-trap where the exclusion window was set at 45 seconds and multiple charge states of the same ion were allowed.

### SAINT Plot Construction

The significance analysis of interactome (SAINT) score (*21*) was determined via the online interface and plots were constructed using ggplot2 (*32*), excluding negative foldchange values.

### GO Term enrichment analysis and interactome construction

GO Term enrichment was performed using the clusterProfiler package in R (*33*). Proteins that were enriched in Raf1-addicted AsPC1 and MM485 pancreatic and melanoma cells at a SAINT score of SAINT ≥ 0.8 were analyzed using “CC”, “BP”, and “MF” ontologies with a Benjamini-Hochberg corrected pValue cutoff of 0.05. STRING(*34*) was used to identify known protein-protein interactions within Raf1 interactors with SAINT ≥ 0.9 and Cytoscape (*35*) was used. Proteins with known mitochondrial localization were labeled based on literature search and Uniprot.

### Mitochondrial buffer purification

To assess inner-mitochondrial localization of exogenously expressed Raf1-BasuHA constructs in MM485 and CHL-1 cells. Assay carried out according to instructs in Mitochondrial Isolation Kit for Cultured Cells (89874, ThermoFisher). Cells digested for 10 minutes on ice with Proteinase K. Proteinase K quenched with 2 mM phenylmethylsulfonyl fluoride (10837091001, Millipore Sigma) for 10 minutes. Samples resuspended in 1X LDS (NP007, NuPage) containing 20 mM DTT (10197777001, Millipore Sigma).

### Mito-tag antibody-based mitochondrial purification

Cells infected with Lentiviral construct containing Mito-tag construct (*23*) and selected in blasticidin for 5 days. Grew appropriate cells to 90% confluence in 15 cm plates. Plates were washed 2X with PBS and aspirated. Cells then scraped into 1 ml of KPBS. Suspension of cells were spun down at 1000 x g for 2 minutes and resuspended in hypotonic lysis buffer RBS (10 mM Tris-HCl, pH 7.4, 3 mM MgCl2, 10 mM NaCl) for MM485 cells, but not Aspc1 cells as they did not require the hypotonic solution for lysis. Cells homogenized with 15 – 20 stokes of PTFE coated homogenizer. Nuclei were cleared with a 1300 x g spin for 5 minutes. Mitochondria-containing supernatant was placed on 200 µl of pre-washed Pierce Anti-HA Magnetic Beads (88837, ThermoFisher). Beads were rotated in 4°C for 3.5 minutes. Mito-bound beads washed 2X 2 minutes with 1 ml KPBS. Mitochondria lysed in 1X RIPA containing Complete Mini Protease Inhibitor Cocktail, EDTA-free (11873580001,Millipore Sigma) PhosStop phosphatase inhibitor cocktail (4906845001,Millipore Sigma). Proceeded to clear lysates and perform western blots.

### Proteinase K digestion of purified mitochondria

For proteinase K digestion of purified mitochondria, 4 µl of thermolabile proteinase K (New England Biolabs, P811S) added to Mitochondria-bound beads resuspended in 30 µl of KPBS. Incubated at 37°C for 8 minutes and placed on ice for an additional 2 minutes. Proteinase K inactivated in a heatblock at 55°C for 10 minutes, with agitation. 15 µl of 4X RIPA added to samples to lyse cells for 10 minutes. Lysate removed and cleared at 16,000 x g before adding LDS (NP0007, NuPage) to 1X with final DTT(10197777001, Millipore Sigma) concentration of 20 mM. Western blots performed.

### Super-resolution image sample preparation

6•10^5^ MM485 cells were plated in a 6-well plate containing a no. 1.5H 170 ± 5 µm thick glass coverslip. Samples were fixed with 4% PFA in PBS for 10 minutes. Samples washed 2x5 minutes with PBS before blocking and permeabilization in 5% BSA, 0.2% Triton X-100 in PBS blocking buffer for 30 minutes. Samples treated with primary mouse Raf1 (12552, Cell Signaling Technology) or Rabbit TOMM20 (ab78547, Abcam) antibodies overnight at 1:250 concentration. Goat anti-Rabbit IgG STAR ORANGE (STORANGE-1002, Abberior) and Goat anti-Mouse IgG STAR RED (STRED-1001) at 1:200 dilution in blocking buffer for 1 hour. Samples further washed 2X with PBS at RT and mounted with ProLong Gold Antifade Mountant (P36930, ThermoFisher).

### Super-resolution image acquisition and analysis

Two-color confocal and 3D STED imaging of primary Mouse Raf1 (12552, Cell Signaling Technology) with Abberior STAR Red antiMouse and rabbit TOMM20 (ab78547) with antiAbberior STAR Orange was performed on an Abberior Facility Line STED microscope. 640 nm and 561 nm excitation was used for Abberior STAR Red and Abberior Star Orange, respectively, while 775 nm depletion was used to achieve super-resolution in both color channels. The focus of the depletion beam was shaped into a ring with zero-intensity in the center, and two high intensity lobes above and below the focal plane for resolution improvement in all three dimensions. Composite images were created and rendered in the PYthon Microscopy Environment (PYME, version >= 23.05.17). Image stack side-views (XZ and YZ) are cropped laterally and then mean-projected along X or Y.

### siRNA nucleofection

Approximately 1•10^6^ cells were mixed per 1.5 nmol siRNA. Cell and siRNA mixture resuspended in 100 µl of Nucleofector Solution for Human Keratinocytes provided with Human Keratinocyte Nucleofector kit (VPD-1002, Lonza). Nucleofection performed on program X-001 of Amaxa Biosystems Nucleofector II electroporator transfection unit. After nucleofection, cells placed in appropriate growth media and then plated and allowed to recover overnight before experiments conducted. Raf1 was targeted using ON-TARGETplus siRNA Raf1(L-0003601-00-0010, Horizon Discovery), GLS was targeted using ON-TARGETplus siRNA GLS (L-004548-01-0010, Horizon Discovery), and ON-TARGETplus nontargeting control was used as a negative control (D-001810-01-20).

### Isotope tracing and harvest

6 x10^5^ MM485 or 4x10^5^ CHL1 cells for each condition used cells plated in a 6-well plate. Cells were incubated for 4 hours in glutamine-free culture media supplemented with 10% dialyzed fetal bovine serum. Afterward, the media was removed and replaced with glutamine-free media supplemented with 10% dFBS and 4 mmol/L U-^13^C_5_-L-glutamine (CLM-1822-H, Cambridge Isotope Laboratories) for 2 hours prior to harvest (n = 3 per condition). Cells were washed twice with ice cold PBS and lysed with 400 µl of acetonitrile in H_2_O. Cells were scraped and sonicated for 30s with a Bioruptor300 sonicator (Diagenode) and spun down at 1.5 x 10^4^ rpm for 10 min at 4°C. 200 µl of supernatant was removed for immediate LC/electrospray ionization MS/MS analysis.

### Metabolite measurement

Quantitative LC/electrospray ionization MS/MS analysis of cell extracts was performed using an Agilent 1290 UHPLC system equipped with an Agilent 6545 quadrupole time-of-flight mass spectrometer. A hydrophilic interaction chromatography method with a BEH amide column (100 × 2.1 mm internal diameter, 1.7 µm; Waters) was used for compound separation at 35°C with a flow rate of 0.3 ml/min. Mobile phase A consisted of 25 mM ammonium acetate and 25 mM ammonium hydroxide in water, and mobile phase B was acetonitrile. The gradient elution was 0– 1 min, 85% B; 1–12 min, 85% B → 65% B; 12–12.2 min, 65% B → 40% B; 12.2–15 min, 40% After the gradient, the column was re-equilibrated at 85% B for 5 min. The overall runtime was 20 min, and the injection volume was 5 µl. Agilent quadrupole time-of-flight was operated in negative mode, and the relevant parameters were ion spray voltage, 3,500 V; nozzle voltage, 1,000 V; fragmentor voltage, 125 V; drying gas flow, 11 liter/min; capillary temperature, 325°C; drying gas temperature, 350°C; and nebulizer pressure, 40 psi. A full scan range was set at 50 to 1,600 m/z. The reference masses were 119.0363 and 980.0164. The acquisition rate was 2 spectra/s. Peak extraction was performed in an Agilent Profinder B.08.00 (Agilent Technologies). The retention time of each metabolite was determined by authentic standards. The mass tolerance was set to ±15 ppm, and retention time tolerance was ±0.2 min. For normalization of ion counts, cell pellets were vacuum dried, and then protein concentration was determined using the Pierce bicinchoninic acid protein assay kit (Thermo Fisher Scientific), according to the manufacturer’s instructions.

### Glutamine/Glutamate Glo glutamine conversion assay

Forty-Thousand cells were plated appropriately and allowed to grow for 24 hours. Cells glutamine starved for approximately 3 hours and glutamine-containing media added to cells for 1.5 hours before lysis. Cells were lysed in 15 µl of 0.3 Normal HCl containing 0.25% DTAB and 30 µl of HCl was immediately added to each well. Plates scraped lightly to dislodge cells after being shaked at 800 RPM for 5 minutes. HCl quenched with 15 µl of 800 mM Tris pH 8.0 and was mixed for 30 seconds. Instructions from Glutamine/Glutamate-Glo Kit (J8021, Promega) were followed past this point.

### Phosphorylation prediction

The group-based prediction system web server was used(*36*). Sequences for the 19 Raf1 interacting proteins were uploaded and the score for the highest-scoring amino acid residue relative to the threshold was provided.

### Immunofluorescence imaging

8 •10^4^ cells grown in 8-well Lab-TEK II glass chamber slides overnight (177380, Thermofisher). Cells were fixed with 4% PFA in PBS for 10 minutes. After fixation, cells washed 2X with PBS and then permeabilized and blocked with 5% BSA containing 0.2% Triton X-100 for 1 hour at RT. Cells washed 2X with PBS and primary antibodies diluted in PBS. Rabbit Glutaminase polyclonal antibody (93434, Abcam) and mouse Raf1 monoclonal antibody (R2404, Millipore Sigma) were diluted 1:50 and secondary Goat anti-Mouse IgG conjugated to Alexa Fluor 488 (A28175, ThermoFisher) or Goat anti-Rabbit IgG conjugated to Alexa Fluor 488 (A-11008, ThermoFisher) at 1:500 dilutions.

### Immunofluorescence quantitation via corrected total cell fluorescence (CTCF)

FIJI (*37*) were used to calculate corrected total cell fluorescence using image J to isolate regions of interest and determine variables to calculate CTCF based on below equation. CTCF = integrated density – (area of selected cell x mean fluorescence of background)

### Co-immunoprecipitation

293T cells were grown in 10 cm plates and transfected with 10 µg of plasmid. Media was changed 16 hours after transfection, and cells were collected 36 hours after transfection. Cell pellets were lysed in 500uL of IP lysis buffer (25 mM Tris-HCl pH 7.4, 150 mM NaCl, 1 mM EDTA, 1% NP-40 and 5% glycerol, complete mini protease inhibitor, PhosStop), briefly sonicated, and cleared by centrifugation at 15000 x g for 10 min at 4°C. For Flag IP, 200 µL of Anti-Flag M2 Slurry (Sigma) were used per sample. For HA IP, 25uL of Anti-HA Magnetic Bead Slurry (Pierce) were used per sample. Beads were incubated with lysates for 2 hours at 4C prior to three washes with lysis buffer. Bound protein was eluted from beads with 1x LDS + 10% BME.

### Recombinant protein purification

Recombinant proteins fused with either Flag or Flag-HA-His tags were transfected at 20 µg per 15 cm plate using Polyethyleneimine 25000 (23966-100, Polysciences) solution at 2.5 µg per 1 µg of DNA. Cells harvested 48 hours post-transfection and lysed on ice for 30 minutes (50 mM Tris-HCl pH 7.5, 300 mM NaCl, 1 mM EDTA, 1% Triton X-100, 1X Protease Inhibitor cocktail (11836170001, MilliporeSigma) and sonicated 3x 10 seconds at 10% amplitude, with pause of at least 10 seconds. The lysate was centrifuged at 16,000 x g for 12 minutes to clear insoluble components and quantified using BCA assay kit. Quantified lysate added to an appropriate amount of anti-FLAG M2 affinity gel (A2220, MilliporeSigma). IP was performed overnight at 4°C with samples rotating head-over-head. Supernatant removed from beads and beads washed 3X 5 minutes with wash buffer (50 mM Tris-HCl pH 7.5, 3 mM EDTA, 0.5% NP-40, 500 mM NaCl, 10% Glycerol, 100 µM DTT). M2 beads were equilibrated with two washes of PBS, and then purified protein was eluted with 3X Flag peptide in PBS at a concentration of 0.5 mg/ml. Elution was performed at 1.5x bead volume for one hour. Eluate concentrated using Amicon 3K MWCO filter columns (MCP003C46, Pall). Proteins quantified using BSA standard curve run alongside the protein on a Bis-Tris gel and stained using InstantBlue Coomassie Protein Stain (ab119211, Abcam).

### Microscale thermophoresis experiments

In-house purified Raf1-Flag-HA-6XHis in 1X PBS diluted to 200 nM in 100 µl. 90 µl of 200 nM protein mixed 1:1 with 100 nM RED-tris-NTA 2^nd^ generation dye (MO-L018, Nanotemper Technologies) and incubated at room temperature for 30 minutes. Labeled samples centrifuged for 10 minutes at 4°C at 15,000 x g and transferred to a fresh tube. Ligand diluted 2X 15 times for a total of 16 tubes. For GLS and MEK2 the highest ligand concentrations were 2 µM and 2.5 µM respectively. Target protein added to samples at a concentration of ∼50 nM and incubated for 5 minutes at RT. Samples placed in Monolith NT.115 capillaries or NT.115 premium capillaries (MO-K022, MO-K-025, Nanotemper Technologies) and measured on Monolith NT.115 instrument (Nanotemper Technologies). Data from 3 independently prepared experiments were analyzed.

### BS3 crosslinking

Raf1 and GLS recombinant protein (50 ng/µl) were crosslinked with either 50, 80, or 120 µM of BS3 (#21580, ThermoFisher) at room temperature for 1 hour and then quenched by adding LDS sample loading buffer. Crosslinked proteins were then run on a gel and subject to in-gel digestion.

### Crosslinked peptide data analysis and circos plot construction

Raw mass spectral data were analyzed using Byonic (Protein Metrics, San Carlos, CA, v2.14.27) to assign peptides and infer proteins. Peptide data were restricted based on tryptic digestion, allowing for n-terminal ragged cleavages, and up to two missed cleavage sites. Proteins were held to a 1% false discovery rate. Byonic X-Link functionality was used to generate predicted crosslinks between biding partners. Data were further analyzed using Byologic (Protein Metrics, San Carlos, CA) for validation, visualization, and report generation. Potential crosslinks were graded empirically based on their log probabilities, their XIC, MS1, and MS/MS spectra, as well as other qualitative features such as the presence of potential collating peptides. The crosslinked peptides were assigned a confidence ranking. Circos plot was made using cx-circos (cx-circos.vercel.app)

### Molecular docking

The structure for RAF1 (Uniprot: P04049) was obtained from PDB, PDB ID: 3OMV (*38*) and the structure for GLSK (UniProt: O94925) was obtained from AlphaFold DB (*39*). For RAF1, Chain A was used as the receptor molecule and the AlphaFold predicted structure of GLSK with the disordered N-terminus truncated, was used as the ligand molecule. The HDOCK server (*40*) was used to perform protein-protein docking in template-free mode. The docking predictions were filtered to select the model with the best score, from which the three-dimensional structure and binding interface were obtained.

### Cell-titer blue growth assay

7,500 MM485 cells were plated 2 x per condition in a 24 well plate. 6 readings were taken with D0 reading being taken 12 hours after plating. 500 µl of CTB reagent per well was mixed and cells were incubated for 2 hours. Assay run in technical triplicate following manufacturer instructions (G8080, Promega).

### Subcutaneous tumor growth experiments

2•10^6^ MM485 cells expressing Raf1-localization constructs(pBABE WT-Raf1-FHH, pBABE RP2-Raf1-FHH, pBABE COX4-Raf1-FHH, pBABE COX4-Raf1(K375A)-FHH) in a retroviral vector were injected with 1:1 PBS and Matrigel (BD Biosciences) and injected subcutaneously into SHO 474 mice (Charles River). Once palpable, tumors were measured using digital calipers and the tumor volume was calculated using the formula (Length/2)*(Width/2)*(Height/2)*4/3π. Researchers were not blinded to group identity.

### Proximity Ligation Assay (PLA) of tumor tissue microarrays

#### Deparaffinization

Proximity ligation of human cancer tissue microarrays (Skin cancer: NBP2-30229, Multiple cancers: NBP2-30263; Novus Biologicals) was used to analyze Raf-GLS protein-protein interaction in diverse cancer contexts. Slides were first incubated at 65°C for 60min. Next, slides were deparaffinized by a series of xylene and ethanol washes: five times for 4min with xylene and two times each for 2min with 100%, 95%, and 75% ethanol. After slide re-equilibration in MiliQ water for 5min, antigen retrieval was performed by submerging the slides for 20min in antigen retrieval solution (1.8mM Citric Acid and 8.2mM Trisodium Citrate) using a steamer constantly held at 97-99°C, and then further incubated in antigen retrieval solution for 20 min at room temperature. Following three washes for 5min with TBST, peroxidase treatment was performed by adding a drop of 3% H_2_O_2_ diluted in water to each slide and incubating them for 10 min at room temperature. Slides were washed again three times for 5min with TBST.

#### Proximity Ligation Assay (PLA)

Protein-protein interaction was measured by PLA with Duolink In Situ Orange Starter Kit Mouse/Rabbit (DUO92102, Millipore Sigma) according to the manufacturer’s instructions. Anti-Raf1 (mouse, 1:50 dilution) (R2404, Millipore Sigma) and anti-GLS (Rabbit, 1:250 dilution) (93434, Abcam) antibodies were applied for PLA primary antibody incubation. After Duolink PLA probe incubation, ligation, and amplification, PLA samples were imaged by Zeiss LSM880 inverted confocal microscopy (Stanford Cell Sciences Imaging Facility). Images were processed using ImageJ(*37*) to improve visualization by uniformly adjusting the brightness and contrast.

#### Quantitation

CellProfiler (*41*) was used to identify and segment nuclei, cells, and PLA punctae in an automated manner. Slides with high tissue background were excluded due to difficulties in accurate automated segmentation. The number of PLA punctae per cell were calculated per image and reported across images of the same type as labeled by Novus biologicals as well as per image/slide.

---

**Fig. S1.**
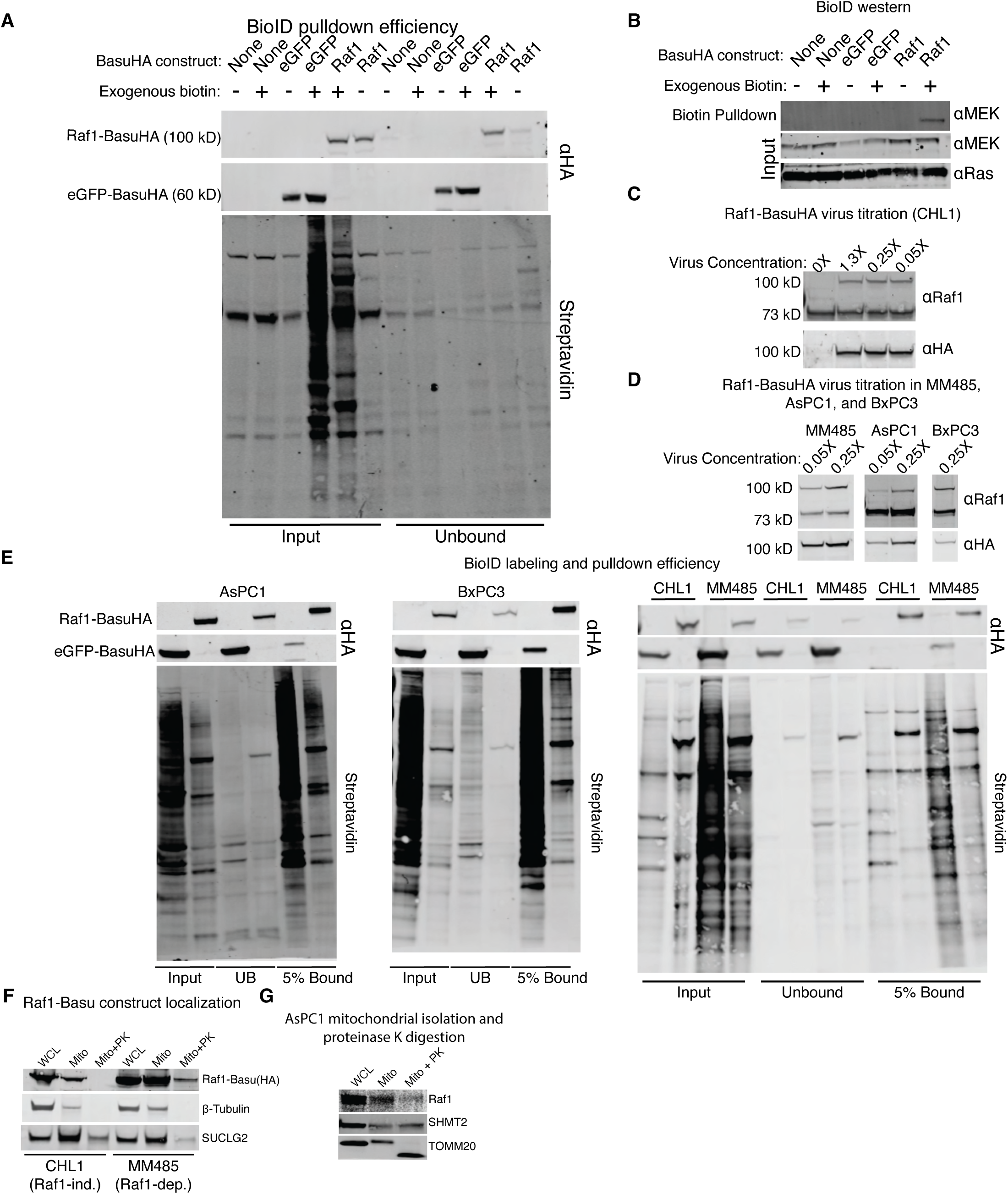
Raf1 proteomics reveals mitochondrial localization. **(A)** Expression of Raf1-BasuHA and eGFP-BasuHA in CHL1 cells. Input is cells post labeling, which occurs with addition of exogenous biotin. Unbound lanes are beads after elution of biotinylated peptides. (**B**) BioID western of eluted proteins blotted for MEK and Ras. (**C**) Concentration of previously concentrated Raf1-BasuHA lentivirus applied to CHL1 cells. Blotted for Raf1 and HA. (**D**) Expression of Raf1-BasuHA in MM485, AsPC1, and BxPC3 cells with varying lentivirus concentration. (**E**) Blots produced from AsPC1, BxPC3, CHL1, and MM485cells expressing Raf1-BasuHA or eGFP-BasuHA with biotin labeling. 5% of streptavidin beads ran on gel for 5% bound. Unbound is running of lysate after elution. (**F**) Buffer-based isolation of mitochondria from MM485 and CHL1 cells and treatment of isolated mitochondria with proteinase K. (**G**) Mito-tag isolation of AsPC1 mitochondria ad blotted for endogenous Raf1.

**Fig. S2.**
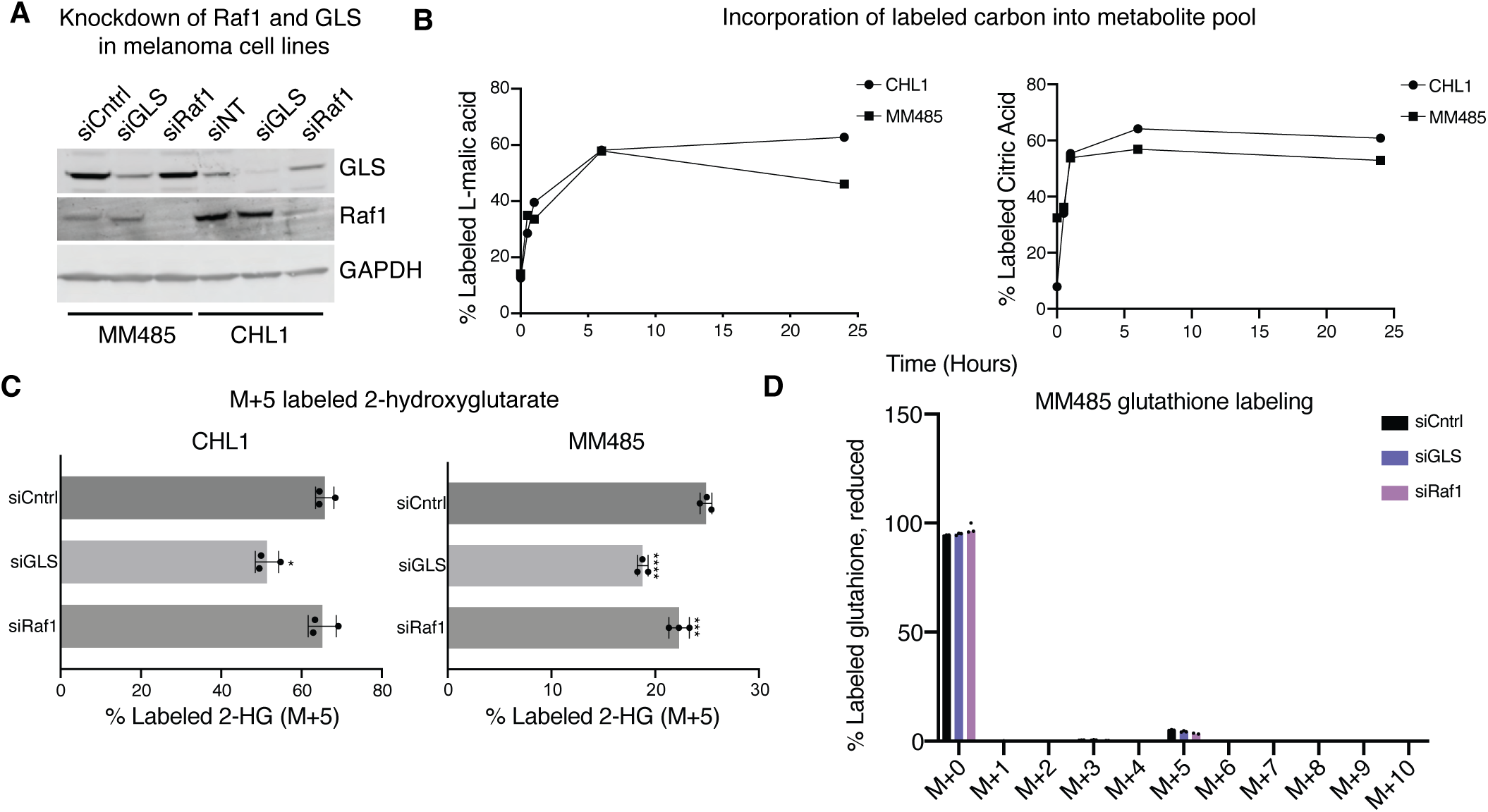
Mitochondrial Raf1 rescues glutamine metabolism. **(A)** siRNA knockdown of Raf1 and GLS in MM485 and CHL1 cells. (**B**) Incorporation of M+5 L-malic acid and L-citric acid at 0 minute, 40 minute, 1 hour, 6 hour, and 24 hour timepoints. Timecourse to determine optimal labeling time. (**C**) Percent labeled 2-HG in CHL1 and MM485 cells; * P < 0.05, *** P < 0.001, **** P < 0.0001. (**D**) Percent labeled isotopologues of reduced glutathione.

**Fig. S3.**
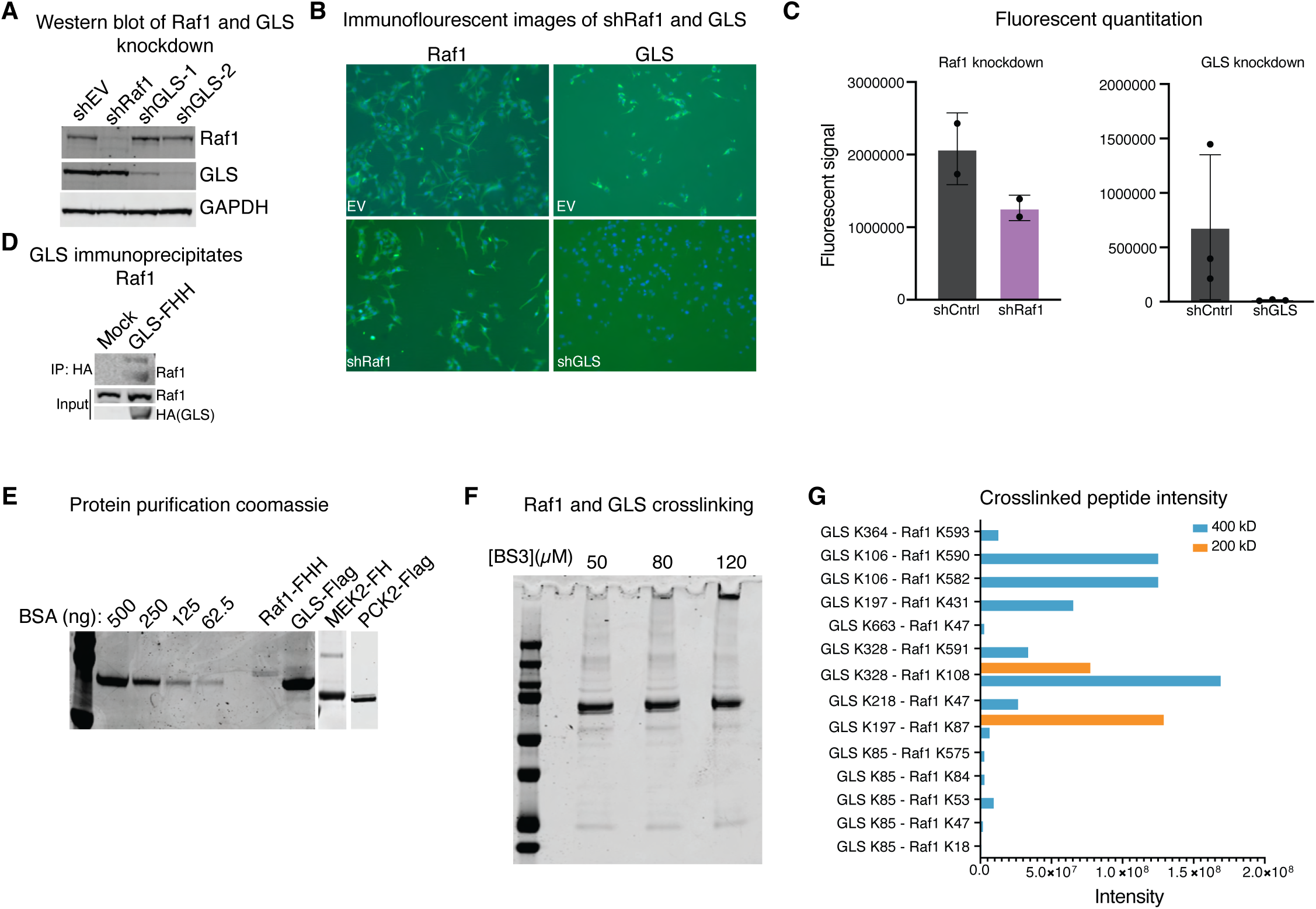
Raf1 directly interacts with GLS. **(A)** shRNA knockdown of Raf1, GLS, or an empty vector (EV) assessed via western blot with GAPDH as a loading control. (**B**) Immunofluorescence of shRNA knockdown of Raf-1, GLS, or an empty vector (EV). Primary antibodies indicated above images. (**C**) Quantitation of immunofluorescence in previous panel via the corrected total cell fluorescence sum across fields of view of several replicates. Presented for both Raf1 and GLS compared to non-targeting short hairpin RNAs (**D**) Immunoprecipitation of GLS-FHH with antiHA antibody followed by immunoblot for Raf1. (**E**) Purification of proteins with a BSA standard curve for in vitro microscale thermophoresis stained with InstantBlue coomassie stain. (**F**) BS3 crosslinking of Raf1 and GLS at three different concentrations of crosslinker. (**G**) Bar graph indicating crosslinks between lysines on GLS and Raf1 by intensity from 200 kD and 400 kD isolated bands.

**Fig. S4.**
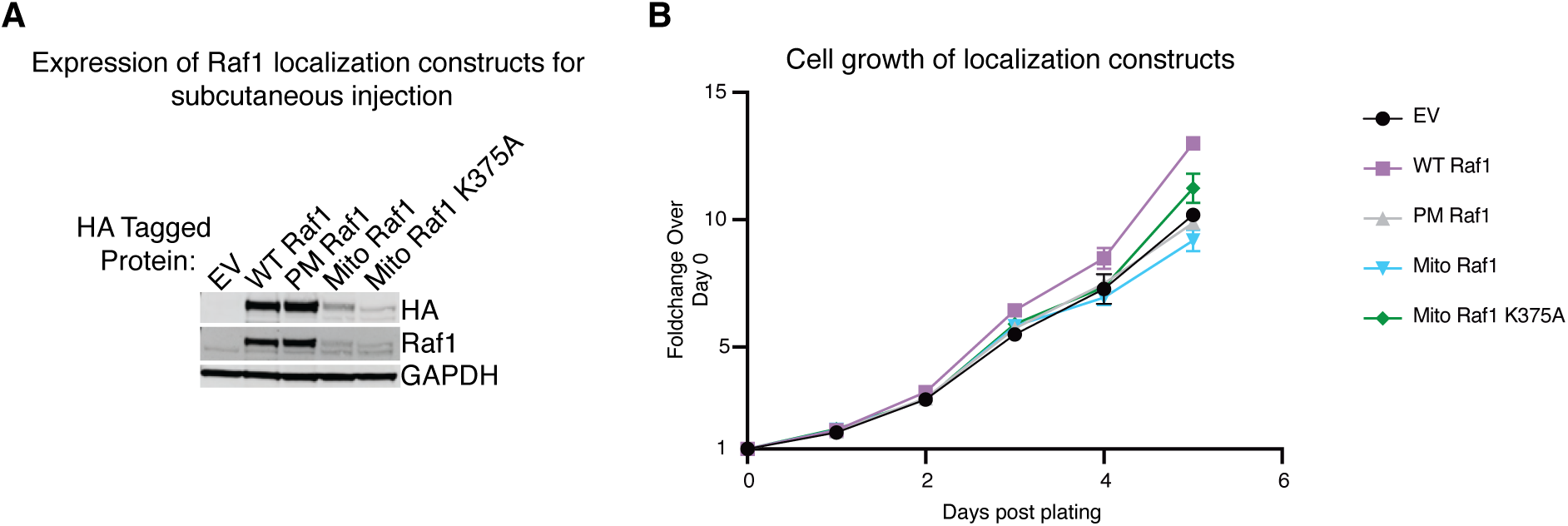
Two-dimensional growth of Raf1 localization constructs. **(A)** Expression of Raf1 WT Raf1, PM Raf1, Mito Raf1, and Mito Raf1 K375A in MM485 cells to be injected subcutaneously in mice. All constructs are FHH tagged and GAPDH used as a loading control. (**B**) Cell-titer blue growth assay measuring cell growth in 2D via fluorescence of localized Raf1 constructs in MM485 cells. Y axis is foldchange in signal over day 0 average for each construct.

**Fig. S5.**
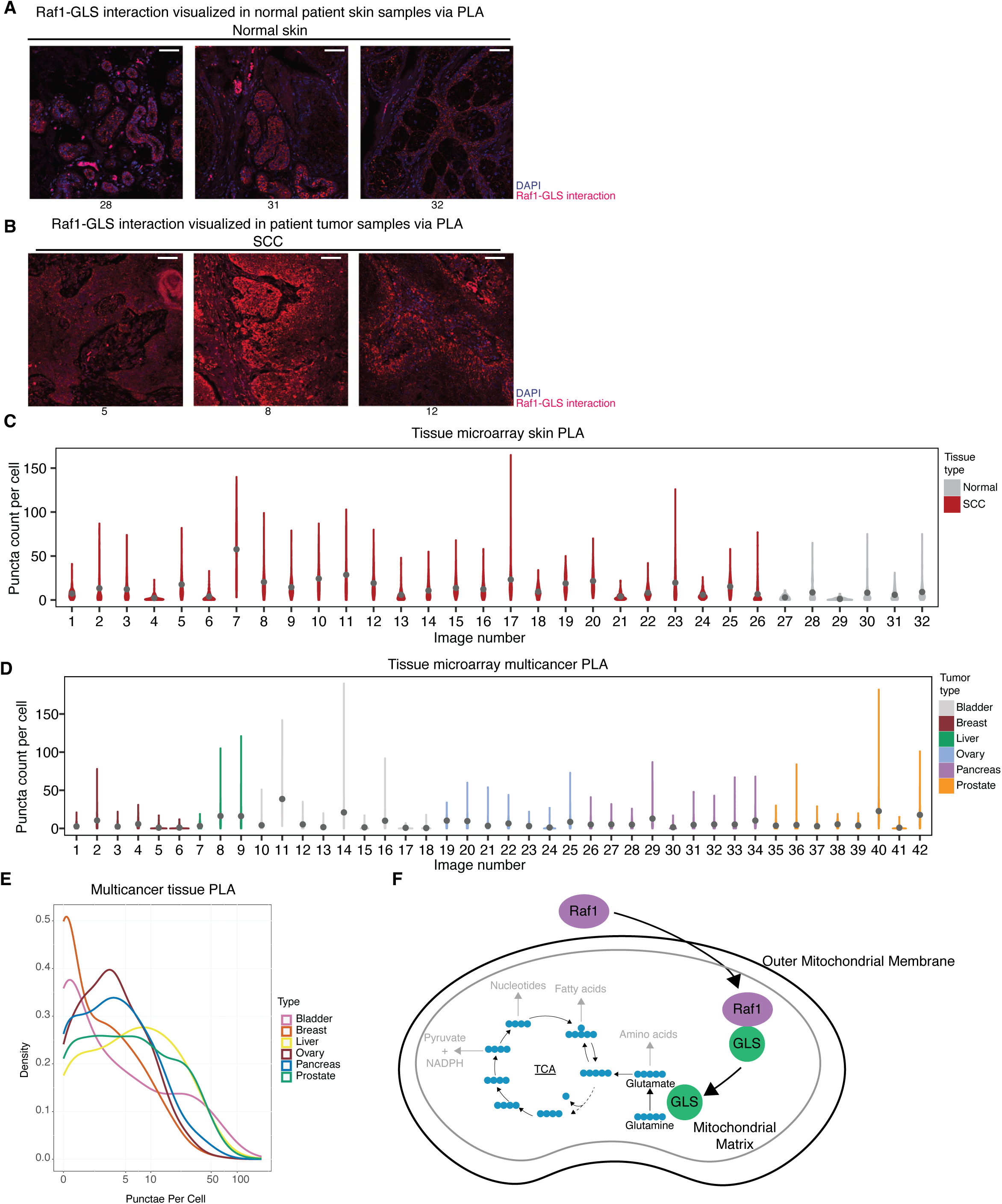
Mitochondrial Raf1 and GLS interaction contributes to tumorigenesis and is present in patient samples. **(A)** Proximity ligation assays for normal patient tissues not depicted previously. PLA signal depicted in red and nuclei stained with DAPI depicted in blue. Numbers correspond to Image Number present in D. (**B**) Proximity ligation assays for 3 squamous cell carcinoma patient samples. Numbers correspond to Image Number present in D. (**C**) Per-image quantitation of PLA punctae per cell in the skin tissue microarray as violin plots. Normal tissues depicted in grey and SCC in red. (**D**) Per-image quantitation of PLA punctae per cell in multicancer tissue microarray. (**E**) Aggregate PLA counts per cell across tissues organized via tumor type presented in a density plot. (**F**) A model of the ways in which Raf1 activation of GLS contributes to tumorigenesis. The black lines have been directly demonstrated by our data, whereas the grey portions are the expected result of Raf1-induced glutaminolysis based on the corpus of literature around glutamine metabolism in cancer.

